# Structure of the proton-powered secretion motor at the heart of the bacterial flagellum

**DOI:** 10.64898/2026.06.13.732081

**Authors:** Mary K. Johnson, Steven Johnson, Justin C. Deme, Luz Alfaro-Alvarado, Owain J. Bryant, Fabienne F.V. Chevance, Kelly T. Hughes, Susan M. Lea

**Affiliations:** Structural Biology, St Jude Children’s Research Hospital, Memphis, TN 38105, USA; Center for Structural Biology, CCR, NCI, Frederick, MD 21702-1201 USA; Department of Biology, University of Utah, Salt Lake City, Utah, USA

## Abstract

Bacterial type III secretion systems export proteins across the inner membrane using the proton-motive force (PMF), but how proton flow is coupled to opening of the export channel is unknown^XX,XX^. Here we present single-particle cryo-EM structures at 2.5-4.2 Å resolution of the intact flagellar Export Apparatus from *Salmonella enterica*, obtained using optimized extraction conditions of the endogenous assembly that retain the complete transmembrane complex. The transmembrane domain of FlhA forms a nonameric funnel-shaped basket beneath the Export Gate, with each subunit harboring a buried pathway of conserved hydrophilic residues spanning the hydrophobic core of the membrane. FlhB is resolved for the first time within the intact gate, revealing helices that thread through the FlhA channel, completely sealing it at rest. A symmetry-free reconstruction captures one FlhA protomer hinged outward at the water-filled cavity, breaking the rotational symmetry. Together, these structures define the architecture of the proton-transducing element of the type III secretion system and suggest a rotary gating mechanism, with parallels to the F_0_ motor of ATP synthase, in which PMF-driven conformational cycling of FlhA drives regulated opening of the export channel.

## Main

The bacterial flagellum is an extraordinary molecular motor, capable of rotating a microns long extracellular filament at speeds exceeding 1000 Hz^1,2^. Construction of the filament is an exquisitely orchestrated process, following a distinct order of sequential assembly of different sub-structures, starting with an inner membrane proximal rod and proceeding via a universal joint structure (the hook) to the filament itself^3,4^. While the basal body that houses the filament is inserted into the bacterial membrane via the general secretion system, the axial structures are secreted via a type III secretion system (T3SS) built into the basal body structure itself^5,6^. Secretion through the T3SS is remarkably rapid, initially proceeding at rates approaching 10,000 amino acids per second, and more than 20,000 protein subunits must be secreted per basal body^7,8^. This poses an acute biophysical challenge, as proteins must be exported through a channel with an inner diameter of up to 20 Å without allowing uncontrolled inward flow of 1 Å protons, while at the same time utilizing the PMF itself for work^9^.

The T3SS itself is a conserved Export Apparatus (EA) found at the center of both the bacterial flagellum and the injectisome virulence machinery of many pathogenic bacteria, including *Shigella*, *Salmonella*, *Yersinia* and *Vibrio* species^10^. The EA was originally defined as an inner membrane complex composed of five proteins, termed FliP, FliQ, FliR, FlhA and FlhB in the flagellar system^11^, associated with a cytoplasmic ATPase complex (FliH, FliI, FliJ) with structural and sequence homology to the F_1_-ATPase^12^. Recent high-resolution single-particle cryo-electron microscopy (cryoEM) structures of purified basal bodies have revealed that the FliP_5_FliQ_4_FliR_1_ sub-complex, termed the Export Gate (EG), is housed within the MS-ring above the conventional membrane plane, where it forms the first turns of the axial helix that becomes the flagellar filament^13–19^. This means that the channel is not merely the conduit through which the structure is built, but the foundation of the structure itself. In the extracted state the gate is completely sealed at the cytoplasmic side by multiple closure points, and no obvious proton pathway exists through the EG that would explain the established role of the PMF^9,20–23^.

The protein channel through the EG must be opened and closed in a highly regulated manner, but the two components most likely to explain this, FlhB and FlhA, have been missing from every high-resolution basal body structure to date (Figure 1a). Our earlier work demonstrated that *Vibrio mimicus* FlhB could be heterologously co-expressed with the FliP_5_FliQ_4_FliR_1_ complex and assemble onto the EG, sitting at the base of the structure with an extended loop sequence forming a lariat structure around the closed base of the EG^14^. The C-terminal domain of FlhB was not observed in this structure, presumably due to the flexibility of the linker connecting it to the hydrophobic region, but crystal structures of this domain are available^24^ and it is implicated in both binding to early secretion substrates via a gate-recognition motif (GRM)^25^, and in the switch to late substrate secretion upon Hook completion^26–28^. Late substrates are proposed to dock to the cytoplasmic domain of FlhA^29,30^, which has been shown to form a nonameric ring structure in several high-resolution studies. However, the proposed TM domain of the FlhA protein has so far proved intractable to high-resolution analysis. Attempts to crystallize the domain alone have proven unsuccessful, cryoEM reconstructions of heterologously expressed full-length protein by multiple groups have only led to structures of the soluble nonamer^31–35^, and basal body extractions have lost the FlhA protein entirely during purification and/or vitrification. Yet the existence of a complex at the critical point between the cytoplasm and the EG has been visualized using low-resolution cryo-electron tomography (cryoET)^36,37^, and genetic evidence has established that the FlhA_TM_ domain is critical for transducing the chemical energy of the PMF into the mechanism of protein export^9,38,39^.

**Figure 1.**
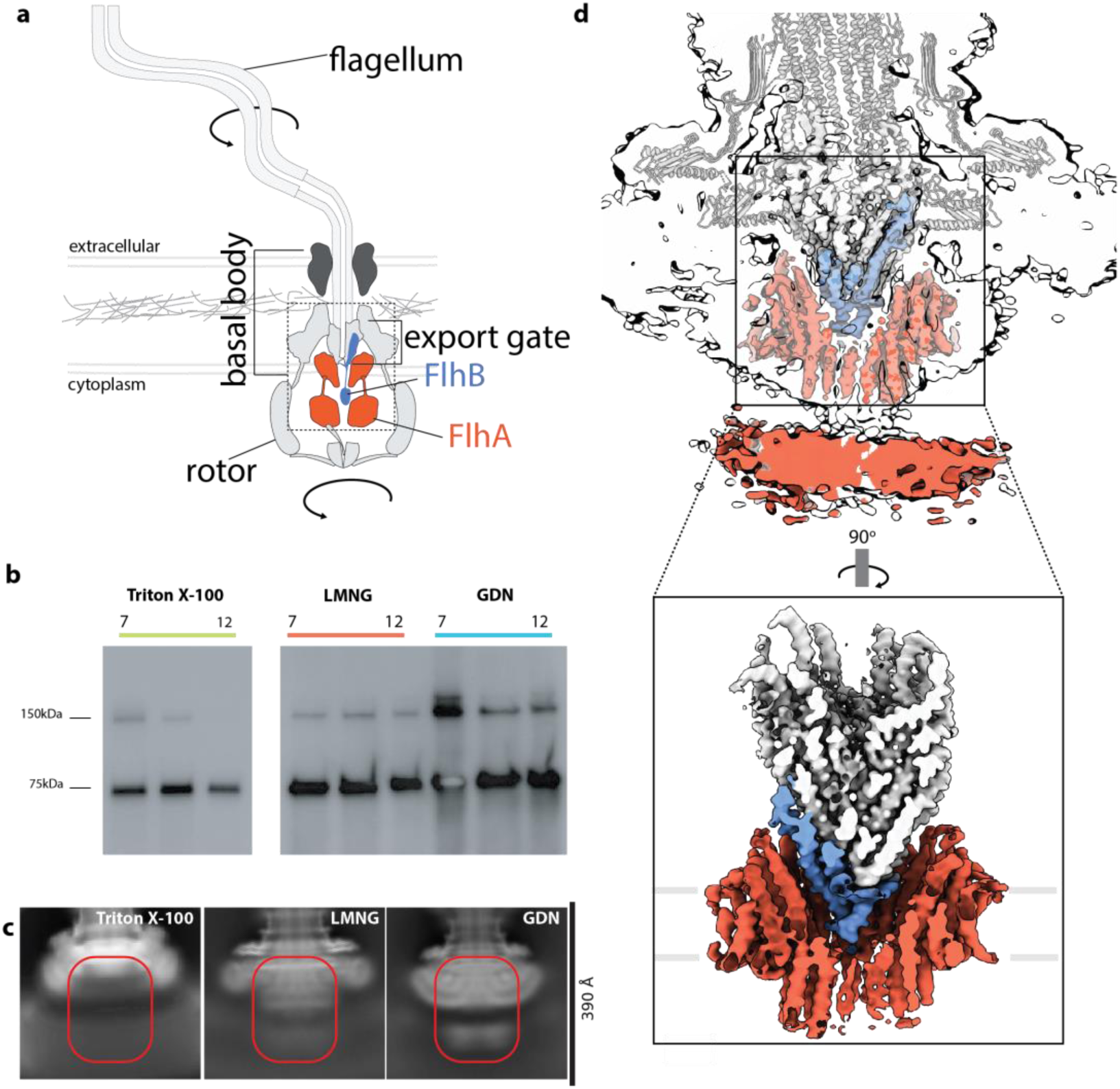
Isolation of the Salmonella Flagellar Basal Body with intact Export Apparatus. **a,** Cartoon of the flagellar assembly across the bacterial membranes highlighting the missing components of the Export Apparatus, FlhB (blue) and FlhA (red). **b,** Western blots from sucrose gradients showing FlhA in the basal body under different detergent extraction conditions. Monomer and dimer bands are observed. **c,** Representative 2D class averages under different extraction conditions. Side views of basal bodies with the Export Apparatus region highlighted. **d,** top, C1 reconstruction of the Salmonella basal body shown at two contour levels and colored as in **(a)**; bottom, zoom on the Export Apparatus with FlhB (blue) and FlhA_TM_ (red). The front two FlhA subunits are hidden for clarity.

Here we present cryoEM structures of the endogenous, intact, flagellar EA from *Salmonella enterica*, obtained by optimizing extraction conditions to retain the full membrane assembly in the context of the intact flagellar basal body, revealing the complex asymmetric architecture of the FlhA_TM_ and previously unseen FlhB regions with respect to the inner membrane and EG. Unexpected features of the full assembly suggest that proton-driven movement of the FlhA_TM_ assembly with respect to the EG may drive protein secretion.

### Structure of the intact *Salmonella* flagellar Export Apparatus

Our earlier structures of native *Salmonella* flagellar basal bodies, extracted in Triton X-100, resolved the FliP_5_FliQ_4_FliR_1_ EG complex at 3.2 Å resolution but lacked density for the FlhA and FlhB components required for secretion^15^. The absence of these components from all single-particle reconstructions to date has left the mechanism by which PMF is coupled to substrate translocation structurally undefined. We therefore screened milder detergent conditions to retain these components during basal body extraction. Western blot analysis of sucrose gradient fractions identified LMNG and GDN as retaining more FlhA in basal body containing fractions (Figure 1b) and samples from both extraction methods were imaged by cryoEM. After particle clean-up, 2D classification revealed distinct micellar structures beneath the position of the EG and evidence of a structure in the cytoplasm consistent with dimensions of the nonameric C-terminal domain of FlhA (Figure 1c). Alignment of the basal body in 3D was performed, producing C1 reconstructions of the rod and EG components to 2.5 Å and 3.0 Å in LMNG and GDN respectively. Both of these volumes contained density for the previously unobserved TM-domain of the FlhB protein, but only the LMNG extracted material resolved protein structure in the region corresponding to the transmembrane region of FlhA (Figure 1d) following extensive classification and local realignments.

The LMNG reconstruction reveals the intact EA within the basal body. Above the inner membrane the EG is housed within the MS-ring, as previously observed, however we now observe FlhB assembled onto the FliP_5_FliQ_4_FliR_1_ structure, completing the *in situ* EG structure. Immediately below the EG, the FlhA TM domain (seen at 3.6 Å) assembles into a funnel shape consistent with position of a cone of density observed in cryoET studies. At lower contours a density extends from the base of the FlhB N-terminal domain and pass through the FlhA basket, while a density consistent in size with the FlhB C-terminal domain is observed to sit on top of the nonameric FlhA cytoplasmic domain.

### The Export Gate is closed in the presence of FlhB

Alignment and classification of the LMNG dataset with a mask around the EG produced a 2.5 Å resolution map with clear density for the entire N-terminal domain of the previously unobserved FlhB (Figure 2a). As in our heterologously expressed FliP_5_FliQ_4_FliR_1_FlhB_1_ structure from *Vibrio mimicus*^14^, FlhB is seen to assemble onto the base of the FliP_5_FliQ_4_FliR_1_ via two long helical hairpins joined by a lariat structure that wraps around the base of the FliQ subunits. Compared to the isolated *Vibrio* EG, the lariat is pulled slightly tighter against the FliQ hairpins above, but otherwise the EG overlays well with the isolated structures and earlier basal body structures. Therefore the EG is observed to be in a closed state in the intact, assembled EA, with FlhB sitting as the lowest point.

**Figure 2.**
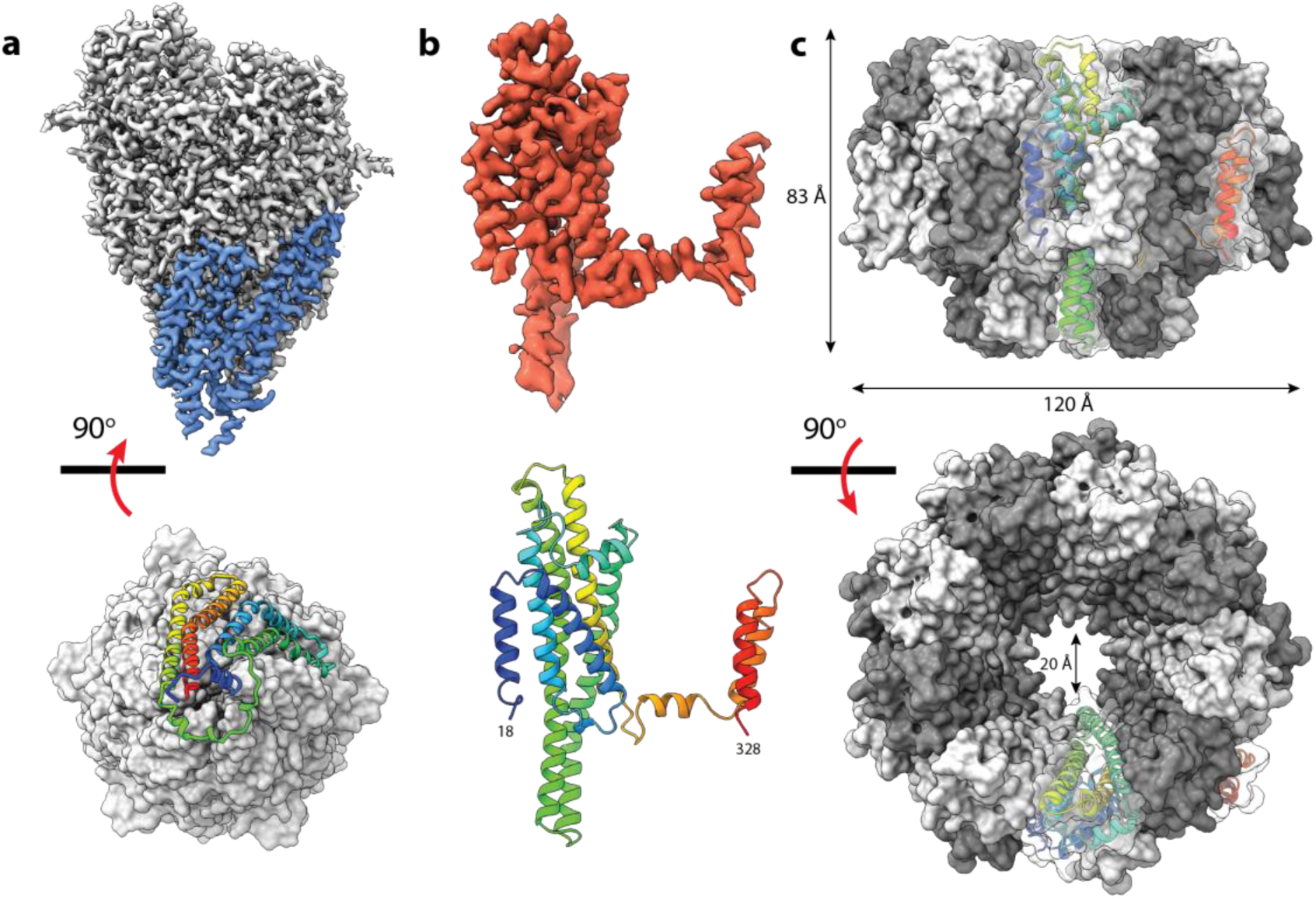
Structure of the Export Apparatus. **a,** CryoEM volume of the Export Gate (gray) with FlhB (blue) residues 9-222 at 2.5 Å. Below is the atomic model viewed from the base of the structure, with the EG represented as a surface and FlhB as cartoon colored N-(blue) to C-terminus (red). **b,** CryoEM volume of a monomer of FlhA (red) residues 18-328 at 3.6 Å, with the built model shown below as a rainbow cartoon. **c,** Structure of the nonamer of FlhA_TM_ with C9 symmetry. One subunit is shown as a rainbow cartoon, while the rest are shown in surface representation with alternating shades of gray to highlight the interwoven nature of the multimer.

### FlhA_TM_ forms a nonameric basket under the Export Gate

An initial C1 reconstruction of the EG centered basal body revealed a clear detergent micelle in the space underneath the EG, containing proteinaceous tubes of density. Classification centered on this region produced classes consistent with the presence of 18 helices in the micelle, consistent with the known nonameric stoichiometry of FlhA. Re-alignment of the particles with the symmetry axis, application of C9 symmetry, and further classification produced maps at 3.6 Å that revealed side-chain level detail and allowed construction and refinement of an atomic model (Figure 2b).

The TM region of FlhA assembles into a nonameric, funnel-shaped basket positioned beneath the EG in the IM. Each monomer adopts an extended conformation which forms extensive contacts with neighboring subunits (Figure 2c). The nonameric ring is built from a compact core formed by TM helices 2-6 at the center, decorated by TM1 from copy n+1 and TMs 7 and 8 from copy n-1 (Figure 2c). The resulting nonamer measures approximately 120 × 80 Å, with a wide periplasmic opening (∼100 Å) that narrows to a ∼20 Å channel at the cytoplasmic face of the membrane. A stretch of residues encompassing the C-terminus of TM4 and the N-terminus of TM5, corresponding approximately to the previously defined FHIPEP domain^40^, projects out of the micelle to create a cytoplasmic cup structure.

### FlhA_TM_ assembly depends on the Export Gate

Surface property mapping (Figure 3a) reveals a hydrophobic belt on the exterior surface of the basket, corresponding to the location of the observed LMNG micelle, thus defining the position of the IM relative to the EA. The inner surface also has a well-defined hydrophobic belt of the right dimensions to interact with the core of the membrane bilayer, but this hydrophobic ring is elevated above the membrane plane defined by the micelle, instead forming a socket that accommodates the hydrophobic exterior of the EG (Figure 3a), despite the symmetry mismatch inherent between the circularly symmetric FlhA nonamer and the pseudo-helical EG assembly.

**Figure 3.**
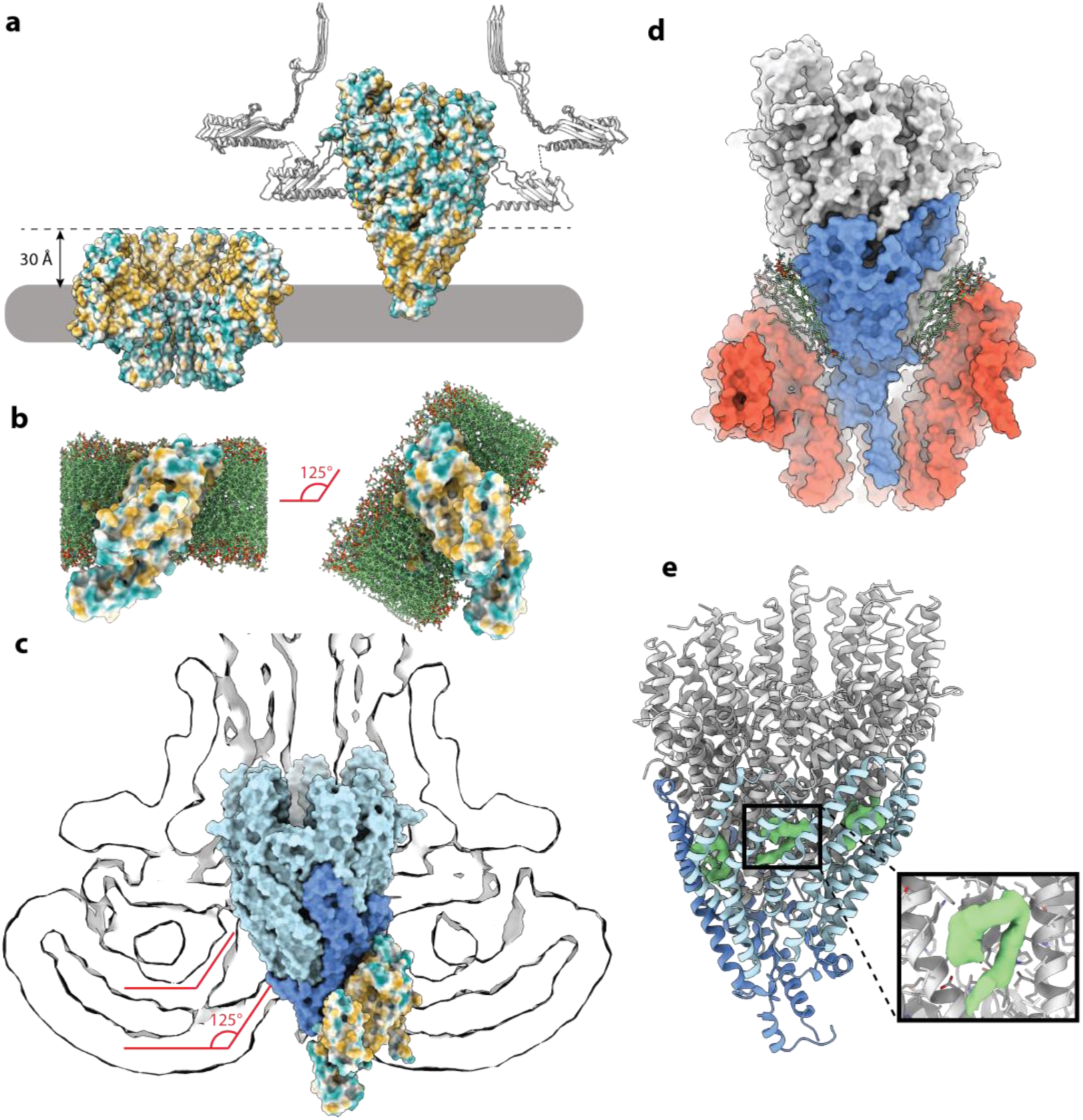
Lipid mediated assembly and function of the Export Apparatus. **a,** Hydrophobic surface analysis of the FlhA_TM_ nonamer demonstrates an external inner membrane facing band and an internal band that sits above the membrane and faces equivalent surfaces on the base of the EG. **b,** FlhA_TM_ monomer embedded in a membrane by MemProtMD (left), and the same simulation reoriented to match the angle of the FlhA_TM_ in the nonameric complex (right). **c,** Slice through a CryoEM reconstruction of a basal body extracted in GDN with no ordered FlhA_TM_ domain, showing the micelle deviating upwards to meet the base of the EG (blue). One copy of FlhA_TM_ is shown with a hydrophobic surface on the right side. **d,** Surface representation of the EG (gray) and FlhB (blue) with the FlhA_TM_ nonamer (red) showing a distinct gap between the two structures. A lipid bilayer is modelled in the gap. **e,** Cartoon representation of the EG and FlhB with unmodelled CryoEM densities sandwiched between the FliP (gray) and FliQ (light blue) monomers. The inset shows a zoom on one such density with the FliQ removed for clarity.

The relative positioning of the internal and external hydrophobic belts has implications for the assembly of FlhA and the wider EA. Molecular simulations indicate that isolated FlhA monomers can be inserted stably in a lipid bilayer (Figure 3b), but in order to satisfy the two major hydrophobic patches on the core they insert at an angle in the membrane that is incompatible with the nonameric assembly. Progressive formation of the nonamer must push the internal hydrophobic band up and out of the membrane layer into the aqueous environment of the periplasm, which would be entropically unfavorable in the absence of the EG to pack against. Therefore the unusual patterning of the hydrophobics on the assembled FlhA nonamer prevents premature pore formation in the absence of the EG to close the structure, preventing formation of the large hydrophilic pore at the center of the FlhATM nonamer (> 20 Å) in the absence of the EG (Figure 2c). However, correct assembly of the EG in the center of the MS ring creates a localized patch of membrane that is pulled upwards to meet the hydrophobic surface at the base of the EG, as previously observed by cryoET and seen in our reconstruction in GDN that lacks ordered FlhA_TM_ nonamer (Figure 3b). The angle of the membrane exactly matches that required to correctly assemble FlhA around the base while satisfying the hydrophobic surfaces.

### Lipids play a key role in the Export Apparatus assembly

Strikingly the hydrophobic belts on the outer surface of the EG and the inner surface of the FlhA are separated by a gap, with no points of direct protein-protein contact (Figure 3c). The dimensions of this gap are sufficient to accommodate a thin lipid bilayer, as modelled by fitting the simulated membrane bound monomer onto the nonamer structure. No lipid densities are observed in the reconstruction, suggesting there are no fixed binding sites on either surface and implying fluidity in that region. Variation in the precise positioning of the FlhA nonamer with respect to the pseudo-helical EG is the likely basis for the need to independently focus on these complexes during alignment to reveal the highest resolution views of each. We also observe novel densities in the EG, sandwiched in a cavity between the FliP and FliQ subunits (Figure 3d), and the completely enclosed and hydrophobic nature of these cavities suggest these densities correspond to trapped lipids. We note that this portion of the EG must undergo movement upon opening of the export aperture, and that the presence of lipids at key interfaces may act as lubricant.

### FlhA_TM_ is a proton channel essential for secretion

Transport of substrates through the EG requires PMF, and the transmembrane region of FlhA has been proposed to be the conduit through which protons flow. The unusual architecture of FlhA_TM_, with an extra layer of lipids interposed between the nonameric basket and the EG above, has direct implications for how this might occur (Figure 4a). The hydrophilic interior of the FHIPEP cup, together with the highly charged residues lining the channel immediately above (including many previously shown to be essential for function), effectively extends the cytoplasmic environment to the closed base of the EG (Figures 4b,c). From this inner ring of essential charged residues, a trail of hydrophilic sidechains can be traced upwards through the hydrophobic core of each monomer to the periplasmic loops of FlhA. The proton path centers on a conserved salt-bridge between R85 and D208, the functional importance of which is well established^38,39^, as mutation disrupts motility and overexpression of a D208A mutant renders membranes leaky to protons^9^. Analysis of the evolutionary conservation of the FlhA demonstrates that all of the highly conserved residues map to the proton path and the inner charge ring (Figure 4d).

**Figure 4.**
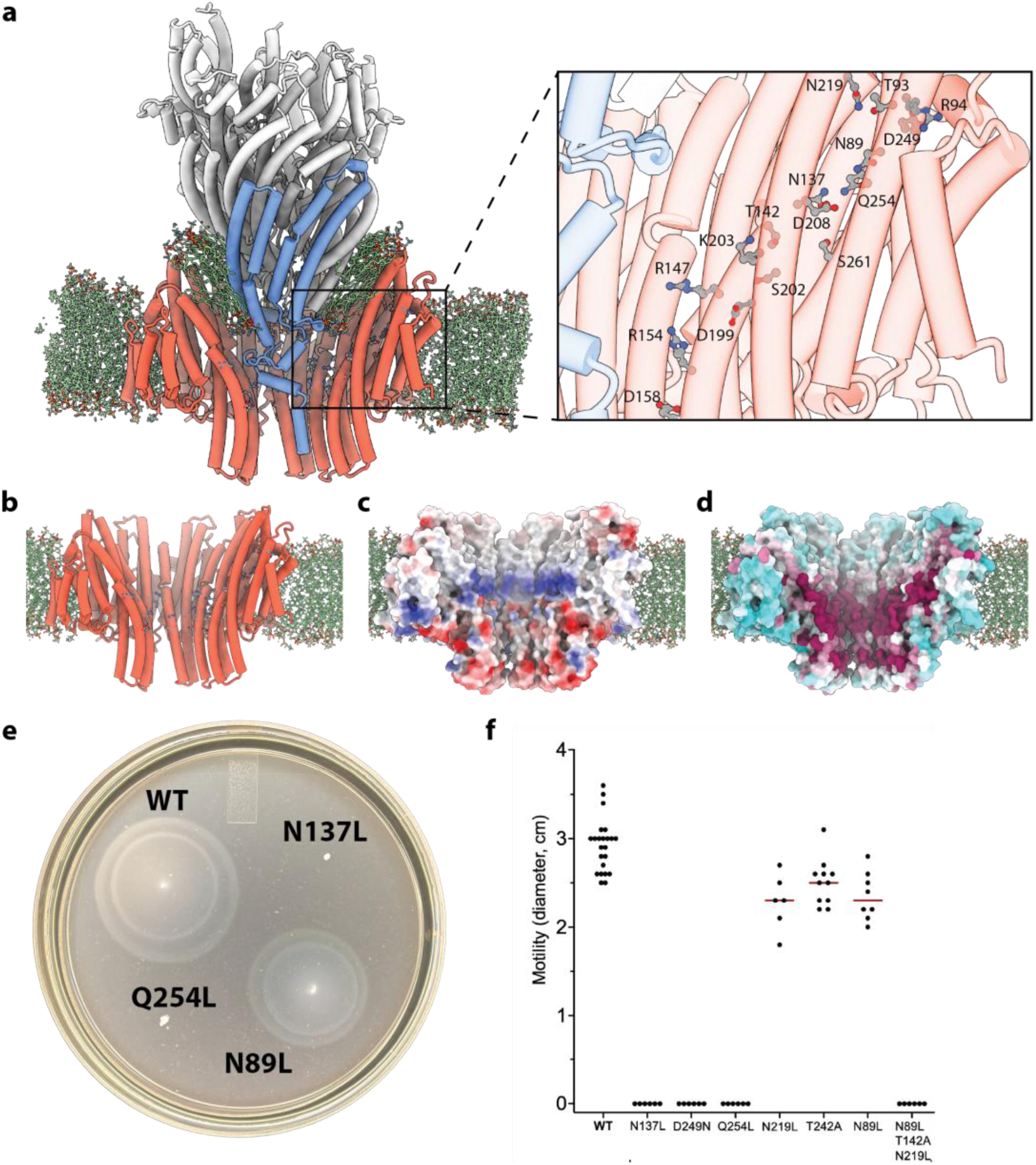
A proton pathway at the heart of the FlhA_TM_ monomer. **a,** Cartoon representation of the EG/B (grey/blue) and FlhA_TM_ nonamer (red) with selected charged sidechains shown on the nonamer, and a hydrophilic pathway through the center of the monomer shown with sidechains. On right is a zoom on the pathway showing key residues. **b,** Cartoon representation of the FlhA_TM_ nonamer showing the concentric rings of charged residues on the inner face. **c,** Electrostatic surface view of the same orientation. **d,** Surface representation colored by sequence conservation shows that the inner charges and proton pathway are the most highly conserved (maroon). **e,** Motility plate assay showing representative strains expressing FlhA protein with point mutations in the hydrophilic pathway. **f,** Graph summarizing motility assays of key point mutants and one triple mutant.

In order to further probe the proton pathway, we made a series of mutations in charged and hydrophilic residues lining the channel. In addition to D208 a negative charge was also required at D249, as previously observed^39^. We also found two non-charged hydrophilic residues to be absolutely essential, N137 and Q254, and identified a further series of hydroxyl and amide containing sidechains that disrupted function when mutated together (Figure 4e,f). These residues define a route from R94 at the top of the funnel to R147 and D199 at the base of the EG, via the D208-R85 salt-bridge. The path of proton conductance is broken in the current model, suggesting it is in a closed state but there are a series of protonatable residues that would require only small movements to form a proton conducting pathway leading to the charged residues in the central channel. Immediately underneath, lining the center of the FlhA ring, are a series of concentric circles of charged residues essential to function (D199, R147, R154, D158)^39^. D158 lines the narrowest constriction point of the channel before a kink around P161 opens the structure back up to form the FHIPEP cytoplasmic cup.

### Asymmetry at the heart of the Export Apparatus

Having defined the folds of FlhB and FlhA within the assembled EA, we reanalyzed the data to examine their structural relationship without any imposed symmetry to try and understand how the closed proton-pathway can be opened and how proton-flow is then coupled to gating of the EG. A subset of LMNG-extracted particles yielded a volume (at 4.2 Å) with functionally relevant additional features (Figure 5a). The N-terminus of FlhB is observed wrapping around a stretch of residues immediately below the transmembrane helices, with co-evolutionary analysis supporting the idea that these contacts reflect a direct physical coupling. The FlhB C-terminal residues then extend beneath this interaction, running nearly parallel to the membrane plane across the mouth of the FlhA basket before turning through 90° and descending through the FHIPEP cup as a helix. This helix does not pass through the center of the channel but is packed against one side. The conformation of these linker helices has implications for the secretion pathway as they occlude the pore in the center of the FlhA_TM_ ring (Figure 5b), adding further closure points to the already closed EG. Docking of AlphaFold model fragments into these densities identifies them as the highly charged helices that form the linker between the FlhB N-and C-terminal domains (Figure 5c) and mutation of clusters of the charged residues are seen to disrupt motility (Figures 5d,e). In addition to the asymmetric positioning of the vertical FlhB linker helix, we observe further asymmetry in the FlhA_TM_ ring (Figure 5f), with one protomer deviating from perfect C9 symmetry, pushed outward by contacts with the FlhB horizontal linker helix.

**Figure 5.**
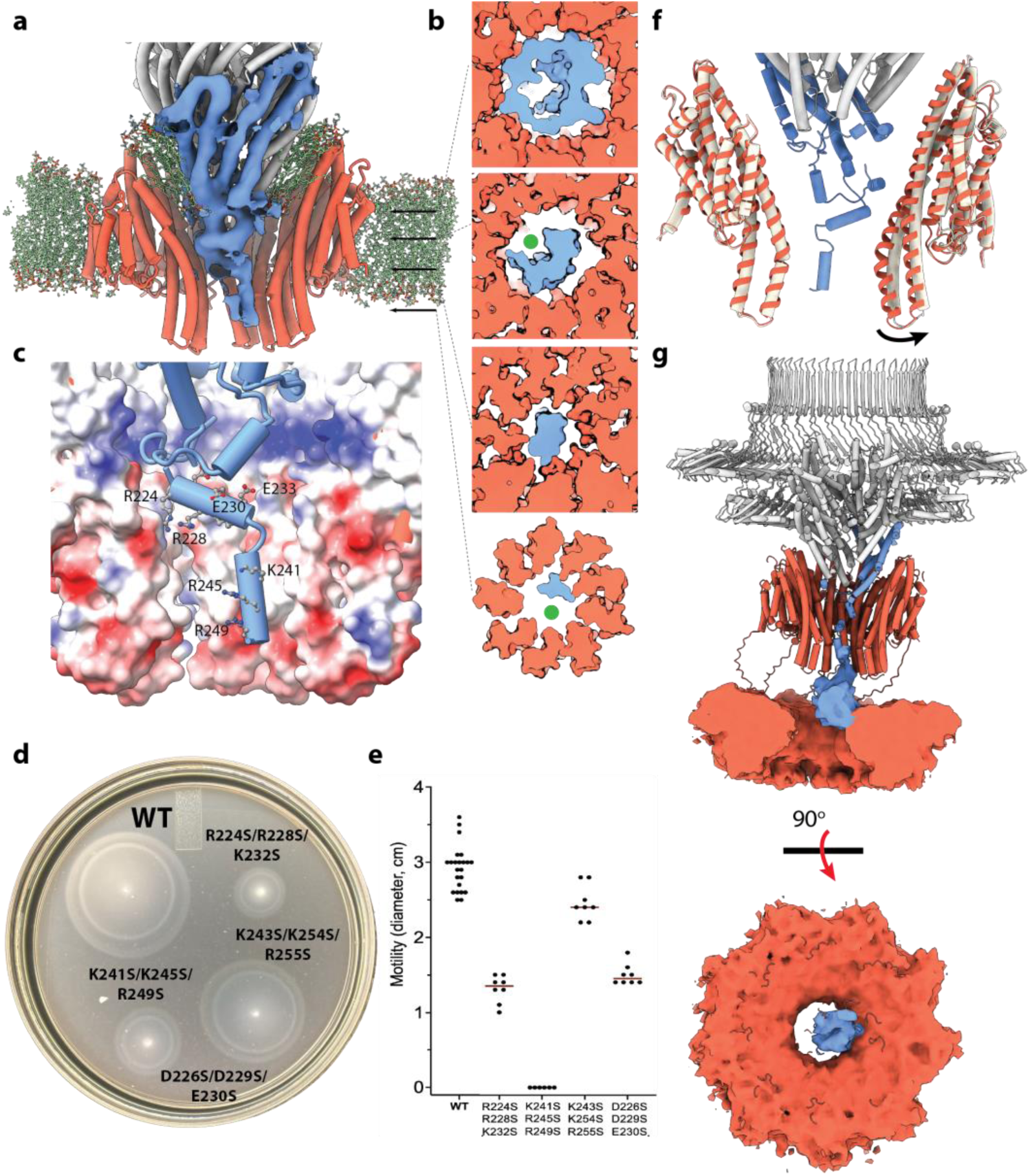
Asymmetry at the heart of the Export Apparatus. **a,** Cartoon representation of the EG (gray) and FlhA_TM_ nonamer (red) with the density for FlhB from a C1 classification showing the FlhB linker extending into the center of the FlhA channel. **b,** Slices through a surface representation of the model built in (a) at different heights in the secretion pathway showing different levels of occlusion of the channel by FlhB. **c,** Zoom on the FlhB linker region in cartoon with key charged sidechains shown as sticks. The inner surface of the FlhA_TM_ nonamer is shown as an electrostatic surface. **d,** Motility plate assay showing representative strains expressing FlhB protein with mutations in the linker. **e,** Graph summarizing motility assays of key triple mutants. **e,** FlhA_TM_ region of the C1 map (light beige) with the cartoon of the C9 model (red) superposed, showing outward movement of a pair of helices in one copy of FlhA_TM_. **g,** Cartoon representation of FliF/EG (gray), FlhB_1-249_ (blue) and FlhA_18-328_ (red) with a low contour map of the cytoplasmic domains of FlhB (blue) and FlhA (red). Below is a top down view of the FlhB_c_/FlhA_c_ complex.

Underneath the FHIPEP cup we observe a lower resolution density that allows docking of the FlhA and FlhB cytoplasmic domains (Figure 5g). The FlhA cytoplasmic nonamer is seen to sit at an angle relative to the symmetry axis of the assembly above and is also displaced laterally. Docked on top of one side of the cytoplasmic ring is a smaller density, consistent in size with the FlhB cytoplasmic domain. This domain sits directly underneath the FHIPEP domain and may drive the displacement of the FlhA cytoplasmic ring.

## Discussion

The flagellar EA sits at the intersection of two fundamental biophysical problems: how to build a micron-scale extracellular organelle through a nanometer-scale channel, and how to harness the proton-motive force to drive that process without catastrophically dissipating the cell’s primary energy currency. Despite decades of genetic and biochemical analysis, the structural basis for PMF coupling to EG opening has remained opaque, largely because the transmembrane components most likely to explain it, FlhA and FlhB, have resisted high-resolution structural characterization in the context of the intact assembly. The structures presented here resolve this impasse, revealing an unexpectedly elaborate transmembrane architecture and suggesting a gating mechanism with striking parallels to rotary proton-translocating motors.

Our structures establish that the transmembrane region of FlhA assembles as a nonameric basket that sits beneath, but is not in direct protein contact with, the EG, separated instead by a lipid-filled interface. This architecture was entirely unanticipated, with prior models assuming intimate protein-protein contacts between FlhA and the FliP₅FliQ₄FliR₁ gate. The lipid layer we observe instead creates a dynamic yet impermeable interface that simultaneously satisfies the hydrophobic requirements of both surfaces while permitting relative movement, analogous to the coupling seen in other proton-driven machines^41–43^. The functional logic is compelling: a rigid connection between FlhA and the EG would prevent the conformational cycling required for gating, while a fluid lipid interface allows PMF-driven movement of the FlhA ring to be transduced into gate opening without requiring a fixed mechanical linkage.

### The intact Export Apparatus defines a new structural paradigm

The assembly logic of the FlhA nonamer also reveals an elegant mechanism for preventing premature pore formation and explains why high level over-expression of FlhA and homologues is easily achieved without promoting cell death^32^. The arrangement of hydrophobic surfaces on the isolated monomer is incompatible with stable, symmetric membrane insertion, as formation of the nonamer drives an internal hydrophobic belt upward and out of the bilayer, creating an energetically unfavorable exposure that is only resolved by docking against the hydrophobic exterior of the EG. The MS-ring housed EG therefore acts as a structural template that both nucleates correct FlhA assembly, via distortion of the inner membrane through an angle that favors the nonamer, and simultaneously closes the large hydrophilic pore at the center of the basket (Figure 6a). This provides a structural rationale for the observed strict dependence of FlhA function on the presence of the assembled EG^44^, and suggests that the sequence of assembly, namely EG first then FlhA, is not merely kinetically convenient but thermodynamically enforced.

**Figure 6.**
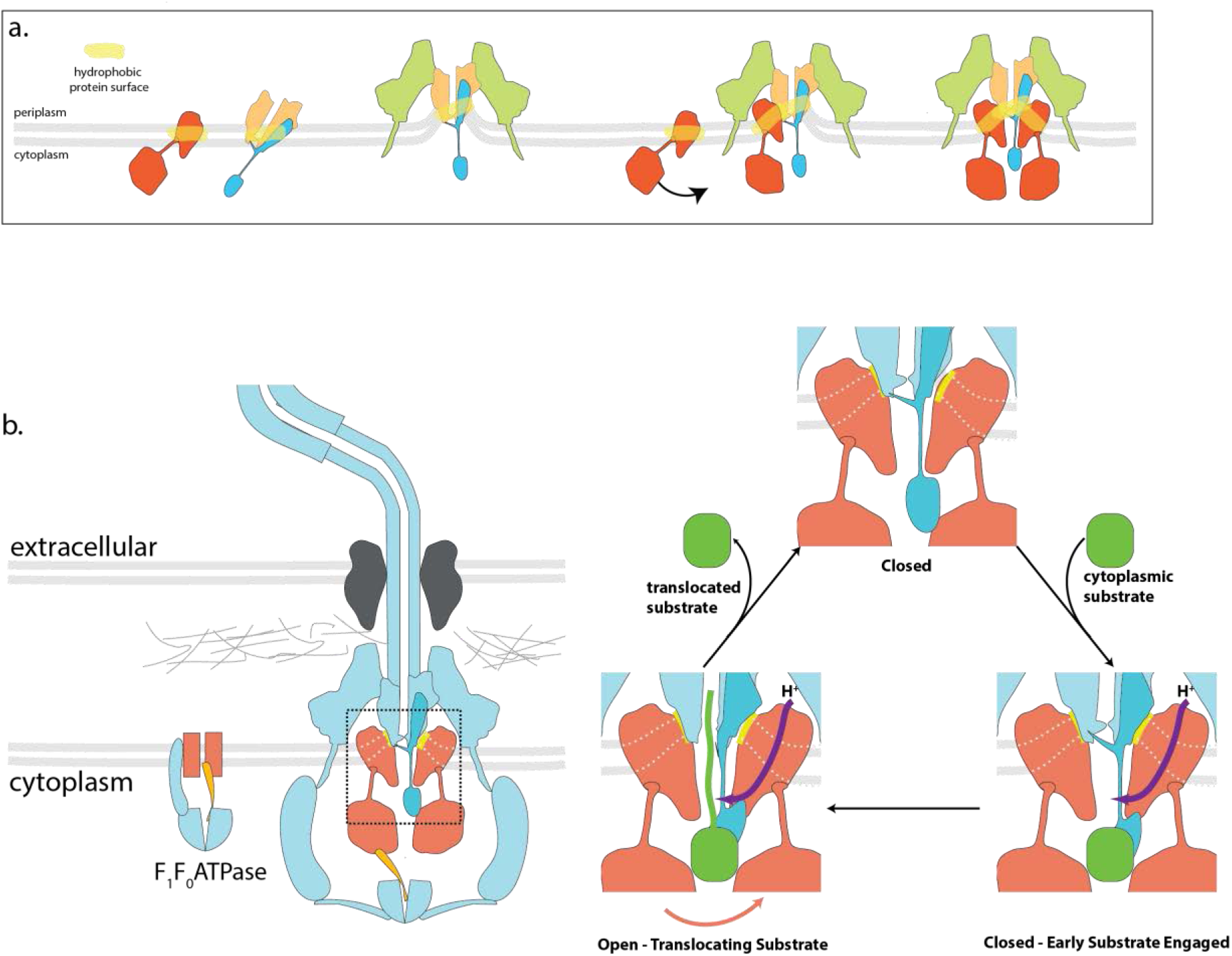
Models of Export Apparatus assembly and function. **a,** Schematic illustrating the role of membrane deformation and the unique hydrophobic patterning of FlhA_TM_ on the regulation of assembly of the Export Apparatus to maintain membrane integrity. **b,** Model showing the relationship of the flagellar Export Apparatus to the F_1_F_O_ ATP Synthase. The assemblies are colored in equivalent fashion to highlight sub-assemblies that are locked together (blue), components that rotate in response to ATP hydrolysis (orange), and membrane embedded proton channels that can rotate (red). The right side shows a potential secretion cycle, with the combination of substrate docking and proton flow through a FlhA_TM_ monomer opening the base of the Export Gate via rotation of the FlhA ring relative to the Export Gate/FlhB complex.

### FlhB as the coupling element

The resolution of FlhB within the intact EA, and particularly the observation of its extended linker region threading through the FlhA basket in the asymmetric reconstruction, identifies FlhB as the structural element that couples conformational changes in FlhA to regulated gate opening. The cluster of FlhB helices immediately below the EG entrance, and the ordered N-terminal peptide of FlhB, are to one side of the structure and positioned to sense or to transmit movements of the FlhA ring. That the linker runs parallel to the membrane plane across the mouth of the basket before descending through the FHIPEP cup places it at precisely the interface where relative FlhA-EG displacement would generate the largest mechanical signal suggesting FlhB is therefore not merely a structural component of the gate but an active transducer. Prior work has established a role for the FlhB C-terminal domain in direct recognition of the early substrates via the so-called GRM motif^25^. The FlhB C-terminal domain, sitting on the upper face of the FlhA cytoplasmic ring as we observe, is well-positioned to present the N-terminus of docked substrate to the EG, transmitting a signal leading to opening of the FlhA proton channel. PMF driven movement of FlhA could then propagate via the asymmetrically positioned FlhB linker to pull on the base of the EG, thereby providing a structural framework for understanding how the substrate recognition and PMF could be coordinated to allow secretion for early substrates (Figure 6b).

### FlhA/FlhB is a proton driven motor

The conserved hydrophilic pathway we identify threading through the hydrophobic core of each FlhA protomer, together with the mutagenesis data defining the essential residues along it, constitutes strong evidence that FlhA is indeed the proton-conducting element of the flagellar T3SS. The pathway is consistent with a Grotthuss-type proton relay^45^, in which proton transfer occurs via sequential protonation and deprotonation of the relay residues rather than by bulk proton diffusion. The identification of N137 and Q254 as absolutely essential non-charged residues is particularly informative since hydroxyl and amide sidechains are established proton relay participants in other systems, including the D-channel of cytochrome c oxidase^46^ and photosynthetic reaction centers^47^, and their requirement here argues for a specific chemical mechanism rather than simple electrostatic guidance. The central R85–D208 salt bridge occupies a critical position in this pathway, and the phenotypes of D208 mutations, particularly the leakiness of cells overexpressing a D208A mutant^9^, are consistent with it playing a key gating role. At the base of the proton pathway, lining the central channel of the FlhA pore, is a cluster of highly conserved charged residues known to being essential for secretion, including R147, R154 and D158.

The architecture we observe, with a lubricated interface between a symmetric proton conducting ring and an asymmetric complex, with highly conserved charges at a single contact point, suggests to us that proton flow likely leads to rotation of FlhA relative to the EG. Such a rotary proton gating mechanism has parallels in the two best characterized rotary motor systems, namely the F_O_ component of the ATP Synthase^41^ and the MotA/B motor of the flagellum itself^42,48^, and also in related rotary motors that regulate other motility^43^ and substrate import events^43,49^. The asymmetry of the EG, when combined with substrate docking at the FlhB C-terminal domain, could open one of the nine proton channels in the FlhA_TM_ ring allowing protons to flow down the PMF gradient and protonate one of the charged residues on the inner surface, triggering a rotation event. In this model, only a single FlhA protomer channel would function as a proton channel at a time, with each being brought sequentially into the activation zone as the wheel turns, and the breaking of the C9 symmetry we observe in one of our structures could be a snapshot of this process. Such a rotation would propagate torsional forces to the base of the EG via FlhB, triggering movements lubricated by the trapped lipids observed in our EG structure.

We also note that the FliH–FliI–FliJ ATPase complex that completes the intact assembly is homologous to the F_1_ component of the ATP Synthase^12,50^ (Figure 6b). The flagellar equivalent of the F_1_ gamma-subunit, FliJ, is essential for efficient utilization of the PMF for protein secretion, and its interaction with the FlhA cytoplasmic domain^51,52^ implicates it in driving rotation of FlhA_cyt_ in response to ATP hydrolysis. Given the direct coupling of the FliI/FliH complex to the EG via the C-ring and MS-ring, this would also drive rotation of FlhA_TM_ ring relative to the EG. In this respect the intact EA would further resemble the ATPase Synthase, consisting of two separate rotor systems, one cytoplasmic and ATP driven, and one membrane localized and PMF driven. How these components interplay to drive maximally efficient secretion remains to be seen, but it is possible the flagellar export machinery represents an evolutionary intermediate between ATP-driven molecular rotary motors and proton-powered secretion systems.

### Conclusions and open questions

The structures presented here provide a high-resolution view of the complete flagellar EA and define the architectural basis for PMF-coupled gating of protein secretion. Several important questions remain: 1) The precise nature of the relative motion between the FlhA ring and the EG. This requires direct experimental interrogation, ideally through time-resolved or substrate-engaged structural studies which are technically extremely challenging and beyond the scope of this manuscript. 2) The mechanism by which hook completion triggers the switch from early to late substrate specificity, in which FlhB plays a central role, remains structurally undefined at the level of the intact EA since our structural information is derived from a pre-switch assembly. Rotation could provide a mechanism for placing the switch proteins under a strain that triggers conformation change or removal of the FlhB_CC_ domain. 3) What role is the FliH–FliI–FliJ rotary ATPase playing, and why can it be dispensed with in the presence of certain permissive mutations. The structural framework established here provides the foundation for addressing each of these questions and for the rational targeting of the EA as an anti-microbial target in flagellated pathogens.

## Methods

### Bacterial strains

Basal bodies were purified from *S.* Typhimurium strain TH25455 (Δ*flgE7659 flhD8070 flhC8092 fliA5225* Δ*fliB-T7771 fliG8835*(ΔPAA) Δ*rflM8403 fljB^enx^ vh2*), which expresses elevated numbers of hook-basal body structures per cell due to elevated FlhDC levels^53^.The *flhD8070*(L22H) and *flhC8092*(Q29P) were isolated as resistant to ClpXP protease degradation^54,55^. The *rflM* gene encodes a repressor of *flhDC* transcription^56^, and was deleted by recombineering using the *tetRA* cassette replacement method. Targeted mutagenesis in *flhA* and *flhB* was conducted using λ-Red recombineering using plasmid helper pSim5 (Cmᴿ). *tetRA* elements derived from transposon Tn10 were first introduced into desired locations in *flhA* or *flhB*, and replaced by the wanted substitutions, as described^57^. PCR or fill-in reactions were conducted using Phusion polymerase with either genomic DNA from *E. coli* strain TH408 (to amplify the *tetRA* cassettes) or from *Salmonella enterica* serovar Typhimurium LT2 strain, or with no template (for fill-in reactions). TetRA elements were introduced into the *Salmonella* genome by selection on tetracycline resistant plates at 37°C while the specific mutations in *flhA* or *flhB* were obtained by selection on fusaric acid plates (12 g/l Apex agar, 5 g/l Bacto tryptone, 5 g/l yeast extract, 10 g/l NaCl, 10 g/l NaH₂PO₄·H₂O, 12 µg/ml fusaric acid, 0.5 µg/ml ATc) at 42°C. Final constructs were confirmed by Sanger sequencing (Eton Biosciences). Produced strains and primers used for are listed in Supplementary Tables S1 and S2.

### Basal body purification

The purification of basal bodies from *S.* Typhimurium strain TH25455 was based on protocols published previously^15^. Briefly, the strain was plated from glycerol stocks on LB agar, then colonies were picked and grown overnight at 37°C in LB medium. Twelve liters of LB medium, in 2.5 L baffled shaker flasks, was inoculated with the overnight culture (13 mL/L) and then incubated at 37°C, 200 rpm until an OD600 of 0.9-1, was reached (approximately 3.5 h). Cells were harvested by centrifugation at 4,000xg for 15 min, 4°C. Cell pellets were resuspended in 240 mL of ice-cold sucrose solution (0.5 M Sucrose, 0.15 M Trizma base (unaltered pH)) and while the cell resuspension was stirred at 4°C, lysozyme and EDTA pH 4.7 were slowly added (over 5 min) to final concentrations of 0.1 mg/ml and 2 mM, respectively. After 5 min of stirring at 4°C, the resuspension was moved to room temperature and stirred slowly for 1 h to allow the formation of spheroplasts. To lyse the cells, 10% LMNG was added to a final concentration of 1% (v/v), and the solution was stirred rapidly for 10 min, until it became translucent. To completely degrade the DNA, 2 mg of DNase I and MgSO4 (5 mM final concentration) were added to the lysate. After 5 min, EDTA pH 4.7 was added to a final concentration of 5 mM. The volume of the lysate was made up to 320 mL with sucrose solution, then unlysed cells and cell debris were removed by centrifugation at 35000xg for 15 min, 4°C. Supernatant was collected and centrifuged at 80,000xg for 50 min, 4°C. Supernatant was collected and centrifuged at 145,000xg for 1 h, 4°C to collect basal bodies. Pellets were resuspended in 1.7 mL of resuspension buffer (0.1 M KCl, 0.3 M Sucrose) with 1% (v/v) LMNG and solubilized in 4°C for 50 mins. Further they were resuspended in 44 mL of resuspension buffer (0.1 M KCl, 0.3 M Sucrose) 0.1% (v/v) LMNG and centrifuged again at 105,000xg for 1 h, 4°C. The pellet was resuspended in 2 mL of HE buffer (10 mM HEPES pH 8, 5 mM EDTA pH 8) with 0.1% (v/v) LMNG, then loaded onto 20-50% (w/w) sucrose gradients in 10 mM HEPES pH 8, 5 mM EDTA pH 8, 0.02% (v/v) LMNG, made with a BioComp Gradient Station, and with 0.2% (v/v) glutaraldehyde added to the 50% sucrose solution. Sucrose gradients were centrifuged for 14 h at 60,000xg, 4°C and then fractionated. Gradient fractions were analyzed by SDS-PAGE and negative stain electron microscopy and those containing basal bodies were pooled and diluted 3-4 times with 10 mM Tris pH 8, 5 mM EDTA pH 8, 0.02% LMNG and centrifuged at 105,000g for 3h at 4°C. Supernatant was removed and the pellet was resuspended in 30μl of 10 mM Tris pH 8, 5 mM EDTA pH 8,0.02% LMNG to prepare the final basal body samples.

### Cryo-EM sample preparation and imaging

Cryo-EM grids were prepared using a Vitrobot Mark IV system (FEI) at a temperature of 4°C and 100% humidity. Basal body samples were applied to Ultrathin Qfoil 300 mesh R 2/1 grids for 60 s before being blotted for 3 s, force -5 and then plunged into liquid ethane. Data were collected in counted mode on a Titan Krios G4 (Thermo Scientific) operating at 300 kV with a Selectris X imaging filter (Thermo Fisher Scientific) with slit width of 10 eV at 165,000x magnification on a Falcon 4 direct detection camera (Thermo Fisher Scientific), with a pixel size was 0.732 Å. Movies were collected at a total dose of 58.5 e^-^/Å ^2^.

### Cryo-EM data processing

Patched motion correction, CTF parameter estimation, particle picking, extraction, and initial 2D classification were performed in SIMPLE 3.0^58,59^. All downstream processing was carried out in cryoSPARC^60^ or RELION^61^,using the csparc2star.py script within UCSF pyem^62^ to convert between formats. Global resolution was estimated from gold-standard Fourier shell correlations (FSCs) using the 0.143 criterion and local resolution estimation was calculated from half maps using phenix.local_resolution^63^. The workflow for image processing is shown in Extended Data Figure 1. Briefly, 189,022 movies were collected and 364,752 particles were obtained after picking and removal of junk classes in SIMPLE and cryoSPARC. An initial subset of the data (34,891 particles) was subjected to multi-class *ab initio* reconstruction (k=2) in cryoSPARC. The resulting good class was lowpass-filtered to 20 Å and used as a reference to align a 232,440 particle subset from the first half of the dataset in non-uniform refinement in C1. Local refinement was carried out using a mask around first the rod and then the EG. Several rounds of 3D classification (cryoSPARC) were carried out with a mask around the EG and classes identified as misalignments due to the pseudosymmetry were realigned and added back to the good alignments. 198,978 remaining particles were subjected to 3D classification using a mask on the micelle region under the EG, producing one class (69,663 particles) with clear density corresponding to helices running through the micelle. Analysis of the micelle region revealed nine-fold symmetry, so the map was aligned to the C9 symmetry axis. The realigned map was lowpass-filtered to 10 Å and used to align the full dataset (364,752 particles) in non-uniform refinement. These particles were Bayesian polished in RELION and used in further focused 3D classification jobs. Classification and local refinement with a mask around the EG/FlhB produced a 2.5 Å map from 151,443 particles. Classification with a mask around both the EG/FlhB and the Flh_TM_ region produced a 4.3 Å map from 58,082 particles. 3D Classification with a mask around the Flh_TM_ was performed producing a class with 19,350 particles. This particle set was expanded using the C9 symmetry and subjected to local refinement, producing a 3.6 Å map. To improve the interpretability of side chains in regions of weaker density EMReady^64^ or deepEMhancer^65^ were used.

### Model building and refinement

Atomic models were built using Coot v0.97 and Coot v1.2^66^. EG coordinates from our earlier basal body structure^15^ (7NVG) were docked into the highest resolution map, followed by docking of an AlphaFold2 model of FlhB. Following rigid body refinement, iterative manual building and real-space refinement into unsharpened, sharpened, deepEMhanced^65^ or EMReady^64^ maps. Final real-space refinement into the B-factor sharpened map with rotamer and Ramachandran restraints was performed in PHENIX^63^. An AlphaFold2^67^ model of residues 18-328 of a FlhA monomer was docked into the C9 expanded map nine times, rigid body refined and iteratively rebuilt and real spaced refined in PHENIX. The EG/FlhB and FlhA_TM_ nonamer models were then combined in the 4.3 Å map, and rigid body and real space refined in PHENIX. Models were validated using Molprobity^68^ within PHENIX. Cryo-EM data collection, image processing and structure refinement statistics are listed in Supplementary Table S3. Figures were prepared using UCSF ChimeraX v.1.12rc202606060206^69^.

### Membrane modelling

Protein models were inserted into bilayers mimicking the *Salmonella enterica* inner membrane using the MemProtMD^70^ Colab Jupyter Notebook: https://colab.research.google.com/github/pstansfeld/MemProtMD/blob/main/MemProtMD_I nsane.ipynb

### Motility Assays

Single colonies from freshly streaked bacteria were inoculated into soft motility agar (10 g tryptone, 5 g NaCl, 3g Difco-Bacto-Agar per liter) by stabbing the plate with a sterile toothpick. Plates were incubated at 37 °C for 4 h 30 min, after which the diameter of the swimming halos was measured. Halo diameters were plotted using GraphPad Prism (version 10). A minimum of six independent colonies were assayed for each strain.

## Data Availability

Cryo-EM volumes and atomic models have been deposited to the EMDB (EMD-77586, EMD-77588, EMD-77603) and PDB (36HU, 36HW, 36IS).

## Code Availability

All code used for cryoEM data analysis, structure determination and refinement are publically available.

## Acknowledgements

We thank Dan Shi (NCI) for assistance with data collection; Hans Elmlund (NCI) for access to SIMPLE code ahead of release. We acknowledge the use of the CCR Center for Structural Biology CryoEM core and the computational resources of the National Institutes of Health (NIH) high performance computing Biowulf cluster. This work was supported by ALSAC (S.M.L.), the Intramural Research Program of the NIH (S.M.L.), and PHS grants R01GM056141 and R56AI184395 from the National Institutes of Health (K.T.H.) The contributions of the NIH author(s) were made as part of their official duties as NIH federal employees, are in compliance with agency policy requirements, and are considered Works of the United States Government. However, the findings and conclusions presented in this paper are those of the author(s) and do not necessarily reflect the views of the NIH or the U.S. Department of Health and Human Services.

## Author Contributions

S.J. and S.M.L. designed the project, interpreted the EM data and built atomic models. M.K.J. optimized the preparation of the basal body samples, made bacterial strains and prepared samples for EM grids. J.C.D made EM grids and together with S.M.L. collected the EM data. O.J.B. made strains and performed secretion experiments. L.A-A. helped with basal body sample preparation. F.F.V.C. and K.T.H. created bacterial strains used for basal body preparation and performed motility studies. S.J. and S.M.L wrote the first draft of the manuscript and all authors commented on manuscript drafts.

---

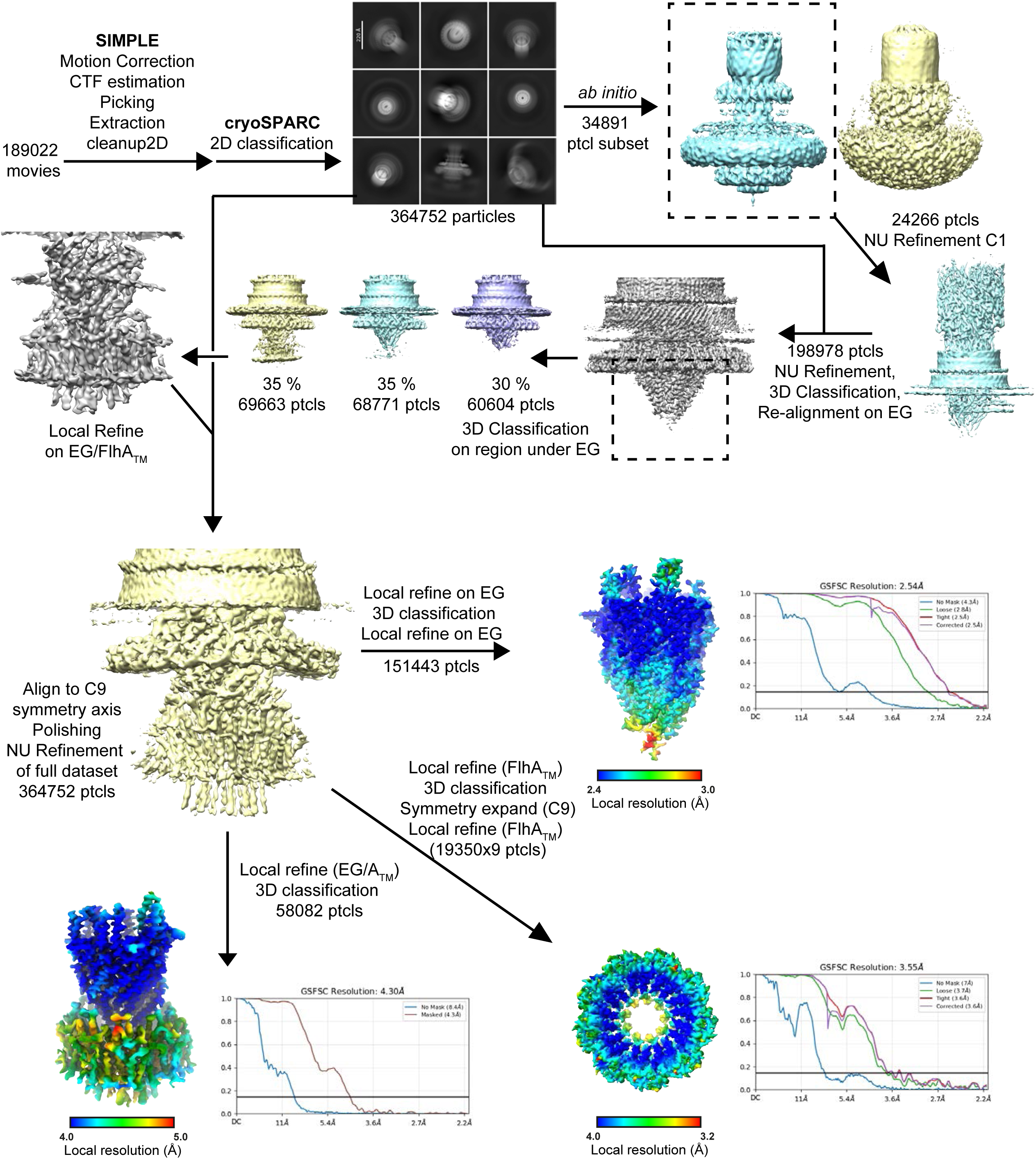

## References

1 Berg, H. C. The Rotary Motor of Bacterial Flagella. Annual Review of Biochemistry 72, 19–54 (2003). 10.1146/annurev.biochem.72.121801.161737

2 Nakamura, S. & Minamino, T. Flagella-Driven Motility of Bacteria. Biomolecules 9 (2019). 10.3390/biom9070279

3 Chevance, F. F. V. & Hughes, K. T. Coordinating assembly of a bacterial macromolecular machine. Nature Reviews Microbiology 6, 455–465 (2008). 10.1038/nrmicro1887

4 Halte, M. & Erhardt, M. Protein Export via the Type III Secretion System of the Bacterial Flagellum. Biomolecules 11 (2021). 10.3390/biom11020186

5 Al-Otaibi, N. S. & Bergeron, J. R. C. in Subcellular Biochemistry 395–420 (2022).

6 Einenkel, R. et al. Building the bacterial flagellum: coordinating regulation, dynamic assembly, and function. Microbiol Mol Biol Rev 89, e0009222 (2025). 10.1128/mmbr.00092-22

7 Halte, M. et al. Bacterial motility depends on a critical flagellum length and energy-optimized assembly. Proc Natl Acad Sci U S A 122, e2413488122 (2025). 10.1073/pnas.2413488122

8 Renault, T. T. et al. Bacterial flagella grow through an injection-diffusion mechanism. Elife 6 (2017). 10.7554/eLife.23136

9 Minamino, T., Morimoto, Y. V., Hara, N., Aldridge, P. D. & Namba, K. The Bacterial Flagellar Type III Export Gate Complex Is a Dual Fuel Engine That Can Use Both H+ and Na+ for Flagellar Protein Export. PLoS Pathog 12, e1005495 (2016). 10.1371/journal.ppat.1005495

10 Worrall, L. J., Majewski, D. D. & Strynadka, N. C. J. Structural Insights into Type III Secretion Systems of the Bacterial Flagellum and Injectisome. Annu Rev Microbiol 77, 669–698 (2023). 10.1146/annurev-micro-032521-025503

11 Minamino, T. & Macnab, R. M. Components of the Salmonella flagellar export apparatus and classification of export substrates. J Bacteriol 181, 1388–1394 (1999). 10.1128/jb.181.5.1388-1394.1999

12 Dreyfus, G., Williams, A. W., Kawagishi, I. & Macnab, R. M. Genetic and biochemical analysis of Salmonella typhimurium FliI, a flagellar protein related to the catalytic subunit of the F0F1 ATPase and to virulence proteins of mammalian and plant pathogens. J Bacteriol 175, 3131–3138 (1993). 10.1128/jb.175.10.3131-3138.1993

13 Kuhlen, L. et al. Structure of the core of the type III secretion system export apparatus. Nat Struct Mol Biol 25, 583–590 (2018). 10.1038/s41594-018-0086-9

14 Kuhlen, L. et al. The substrate specificity switch FlhB assembles onto the export gate to regulate type three secretion. Nat Commun 11, 1296 (2020). 10.1038/s41467-020-15071-9

15 Johnson, S. et al. Molecular structure of the intact bacterial flagellar basal body. Nat Microbiol 6, 712–721 (2021). 10.1038/s41564-021-00895-y

16 Johnson, S. et al. Structural basis of directional switching by the bacterial flagellum. Nat Microbiol 9, 1282–1292 (2024). 10.1038/s41564-024-01630-z

17 Tan, J. et al. Structural basis of assembly and torque transmission of the bacterial flagellar motor. Cell 184, 2665–2679.e2619 (2021). 10.1016/j.cell.2021.03.057

18 Tan, J. et al. Structural basis of the bacterial flagellar motor rotational switching. Cell Res 34, 788–801 (2024). 10.1038/s41422-024-01017-z

19 Singh, P. K. et al. CryoEM structures reveal how the bacterial flagellum rotates and switches direction. Nat Microbiol 9, 1271–1281 (2024). 10.1038/s41564-024-01674-1

20 Paul, K., Erhardt, M., Hirano, T., Blair, D. F. & Hughes, K. T. Energy source of flagellar type III secretion. Nature 451, 489–492 (2008). 10.1038/nature06497

21 Minamino, T., Morimoto, Y. V., Hara, N. & Namba, K. An energy transduction mechanism used in bacterial flagellar type III protein export. Nat Commun 2, 475 (2011). 10.1038/ncomms1488

22 Minamino, T., Morimoto, Y. V., Kinoshita, M. & Namba, K. Membrane voltage-dependent activation mechanism of the bacterial flagellar protein export apparatus. Proc Natl Acad Sci U S A 118 (2021). 10.1073/pnas.2026587118

23 Terashima, H. et al. In Vitro Reconstitution of Functional Type III Protein Export and Insights into Flagellar Assembly. mBio 9 (2018). 10.1128/mBio.00988-18

24 Deane, J. E., Abrusci, P., Johnson, S. & Lea, S. M. Timing is everything: the regulation of type III secretion. Cell Mol Life Sci 67, 1065–1075 (2010). 10.1007/s00018-009-0230-0

25 Bryant, O. J., Dhillon, P., Hughes, C. & Fraser, G. M. Recognition of discrete export signals in early flagellar subunits during bacterial type III secretion. Elife 11 (2022). 10.7554/eLife.66264

26 Chevance, F. F. V. et al. Active removal of inhibitory components drives the flagellar type 3 secretion-specificity switch. mBio, e0103726 (2026). 10.1128/mbio.01037-26

27 Inoue, Y., Kinoshita, M., Namba, K. & Minamino, T. Mutational analysis of the C-terminal cytoplasmic domain of FlhB, a transmembrane component of the flagellar type III protein export apparatus in Salmonella. Genes Cells 24, 408–421 (2019). 10.1111/gtc.12684

28 Minamino, T., Inoue, Y., Kinoshita, M. & Namba, K. FliK-Driven Conformational Rearrangements of FlhA and FlhB Are Required for Export Switching of the Flagellar Protein Export Apparatus. J Bacteriol 202 (2020). 10.1128/JB.00637-19

29 Xing, Q. et al. Structures of chaperone-substrate complexes docked onto the export gate in a type III secretion system. Nat Commun 9, 1773 (2018). 10.1038/s41467-018-04137-4

30 Kinoshita, M. et al. Rearrangements of alpha-helical structures of FlgN chaperone control the binding affinity for its cognate substrates during flagellar type III export. Mol Microbiol 101, 656–670 (2016). 10.1111/mmi.13415

31 Abrusci, P. et al. Architecture of the major component of the type III secretion system export apparatus. Nat Struct Mol Biol 20, 99–104 (2013). 10.1038/nsmb.2452

32 Kuhlen, L., Johnson, S., Cao, J., Deme, J. C. & Lea, S. M. Nonameric structures of the cytoplasmic domain of FlhA and SctV in the context of the full-length protein. PLoS One 16, e0252800 (2021). 10.1371/journal.pone.0252800

33 Matthews-Palmer, T. R. S. et al. Structure of the cytoplasmic domain of SctV (SsaV) from the Salmonella SPI-2 injectisome and implications for a pH sensing mechanism. J Struct Biol 213, 107729 (2021). 10.1016/j.jsb.2021.107729

34 Majewski, D. D., Lyons, B. J. E., Atkinson, C. E. & Strynadka, N. C. J. Cryo-EM analysis of the SctV cytosolic domain from the enteropathogenic E. coli T3SS injectisome. J Struct Biol 212, 107660 (2020). 10.1016/j.jsb.2020.107660

35 Yuan, B. et al. Structural Dynamics of the Functional Nonameric Type III Translocase Export Gate. J Mol Biol 433, 167188 (2021). 10.1016/j.jmb.2021.167188

36 Butan, C., Lara-Tejero, M., Li, W., Liu, J. & Galán, J. E. High-resolution view of the type III secretion export apparatus in situ reveals membrane remodeling and a secretion pathway. Proc Natl Acad Sci U S A 116, 24786–24795 (2019). 10.1073/pnas.1916331116

37 Chen, S. et al. Structural diversity of bacterial flagellar motors. Embo j 30, 2972–2981 (2011). 10.1038/emboj.2011.186

38 Hara, N., Namba, K. & Minamino, T. Genetic characterization of conserved charged residues in the bacterial flagellar type III export protein FlhA. PLoS One 6, e22417 (2011). 10.1371/journal.pone.0022417

39 Erhardt, M. et al. Mechanism of type-III protein secretion: Regulation of FlhA conformation by a functionally critical charged-residue cluster. Mol Microbiol 104, 234–249 (2017). 10.1111/mmi.13623

40 Barker, C. S., Inoue, T., Meshcheryakova, I. V., Kitanobo, S. & Samatey, F. A. Function of the conserved FHIPEP domain of the flagellar type III export apparatus, protein FlhA. Mol Microbiol 100, 278–288 (2016). 10.1111/mmi.13315

41 Mühleip, A., McComas, S. E. & Amunts, A. Structure of a mitochondrial ATP synthase with bound native cardiolipin. Elife 8 (2019). 10.7554/eLife.51179

42 Deme, J. C. et al. Structures of the stator complex that drives rotation of the bacterial flagellum. Nat Microbiol 5, 1553–1564 (2020). 10.1038/s41564-020-0788-8

43 Hennell James, R., et al. Structure and mechanism of the proton-driven motor that powers type 9 secretion and gliding motility. Nat Microbiol 6, 221–233 (2021). 10.1038/s41564-020-00823-6

44 Morimoto, Y. V. et al. Assembly and stoichiometry of FliF and FlhA in Salmonella flagellar basal body. Mol Microbiol 91, 1214–1226 (2014). 10.1111/mmi.12529

45 Agmon, N. The Grotthuss mechanism. Chemical Physics Letters 244, 456–462 (1995). 10.1016/0009-2614(95)00905-J

46 Reidelbach, M. & Imhof, P. Proton transfer in the D-channel of cytochrome c oxidase modeled by a transition network approach. Biochim Biophys Acta Gen Subj 1864, 129614 (2020). 10.1016/j.bbagen.2020.129614

47 Sugo, Y. & Ishikita, H. Mechanism of Asparagine-Mediated Proton Transfer in Photosynthetic Reaction Centers. Biochemistry 62, 1544–1552 (2023). 10.1021/acs.biochem.3c00013

48 Santiveri, M. et al. Structure and Function of Stator Units of the Bacterial Flagellar Motor. Cell 183, 244–257.e216 (2020). 10.1016/j.cell.2020.08.016

49 Celia, H. et al. Cryo-EM structures of the E. coli Ton and Tol motor complexes. Nat Commun 16, 5506 (2025). 10.1038/s41467-025-61286-z

50 Majewski, D. D. et al. Cryo-EM structure of the homohexameric T3SS ATPase-central stalk complex reveals rotary ATPase-like asymmetry. Nat Commun 10, 626 (2019). 10.1038/s41467-019-08477-7

51 Jensen, J. L., Yamini, S., Rietsch, A. & Spiller, B. W. “The structure of the Type III secretion system export gate with CdsO, an ATPase lever arm”. PLoS Pathog 16, e1008923 (2020). 10.1371/journal.ppat.1008923

52 Ibuki, T. et al. Interaction between FliJ and FlhA, components of the bacterial flagellar type III export apparatus. J Bacteriol 195, 466–473 (2013). 10.1128/JB.01711-12

53 Erhardt, M. & Hughes, K. T. C-ring requirement in flagellar type III secretion is bypassed by FlhDC upregulation. Mol Microbiol 75, 376–393 (2010). 10.1111/j.1365-2958.2009.06973.x

54 Sato, Y., Takaya, A., Mouslim, C., Hughes, K. T. & Yamamoto, T. FliT selectively enhances proteolysis of FlhC subunit in FlhD4C2 complex by an ATP-dependent protease, ClpXP. J Biol Chem 289, 33001–33011 (2014). 10.1074/jbc.M114.593749

55 Takaya, A. et al. YdiV: a dual function protein that targets FlhDC for ClpXP-dependent degradation by promoting release of DNA-bound FlhDC complex. Mol Microbiol 83, 1268–1284 (2012). 10.1111/j.1365-2958.2012.08007.x

56 Singer, H. M., Erhardt, M. & Hughes, K. T. RflM functions as a transcriptional repressor in the autogenous control of the Salmonella Flagellar master operon flhDC. J Bacteriol 195, 4274–4282 (2013). 10.1128/jb.00728-13

57 Karlinsey, J. E. lambda-Red genetic engineering in Salmonella enterica serovar Typhimurium. Methods Enzymol 421, 199–209 (2007). 10.1016/s0076-6879(06)21016-4

58 Caesar, J. et al. SIMPLE 3.0. Stream single-particle cryo-EM analysis in real time. J Struct Biol X 4, 100040 (2020). 10.1016/j.yjsbx.2020.100040

59 Van, C. T. S. et al. Probabilistic single-particle cryo-EM ab initio 3D reconstruction in SIMPLE. Acta Crystallogr D Struct Biol 81, 396–409 (2025). 10.1107/s2059798325005686

60 Punjani, A., Rubinstein, J. L., Fleet, D. J. & Brubaker, M. A. cryoSPARC: algorithms for rapid unsupervised cryo-EM structure determination. Nat Methods 14, 290–296 (2017). 10.1038/nmeth.4169

61 Zivanov, J. et al. New tools for automated high-resolution cryo-EM structure determination in RELION-3. Elife 7 (2018). 10.7554/eLife.42166

62 Asarnow, D., Palovcak, E. & Cheng, Y.

63 Afonine, P. V. et al. Real-space refinement in PHENIX for cryo-EM and crystallography. Acta Crystallogr D Struct Biol 74, 531–544 (2018). 10.1107/s2059798318006551

64 He, J., Li, T. & Huang, S. Y. Improvement of cryo-EM maps by simultaneous local and non-local deep learning. Nat Commun 14, 3217 (2023). 10.1038/s41467-023-39031-1

65 Sanchez-Garcia, R. et al. DeepEMhancer: a deep learning solution for cryo-EM volume post-processing. Commun Biol 4, 874 (2021). 10.1038/s42003-021-02399-1

66 Brown, A. et al. Tools for macromolecular model building and refinement into electron cryo-microscopy reconstructions. Acta Crystallogr D Biol Crystallogr 71, 136–153 (2015). 10.1107/s1399004714021683

67 Jumper, J. et al. Highly accurate protein structure prediction with AlphaFold. Nature 596, 583–589 (2021). 10.1038/s41586-021-03819-2

68 Williams, C. J. et al. MolProbity: More and better reference data for improved all-atom structure validation. Protein Sci 27, 293–315 (2018). 10.1002/pro.3330

69 Meng, E. C. et al. UCSF ChimeraX: Tools for structure building and analysis. Protein Sci 32, e4792 (2023). 10.1002/pro.4792

70 Stansfeld, P. J. et al. MemProtMD: Automated Insertion of Membrane Protein Structures into Explicit Lipid Membranes. Structure 23, 1350–1361 (2015). 10.1016/j.str.2015.05.006

