## Supplementary Tables S1 to S3 for "Structure of the proton-powered secretion motor at the heart of the bacterial flagellum"

**Table S1. List of Strains**

| **Strain**  **Number** | **Genotype** | **Recipient strain** | **Donor**  **(Template strain/primers)** |
| --- | --- | --- | --- |
| LT2 | WT *Salmonella* Typhimurium | KTH strain |  |
| TH27083 | pSim-5/LT2 | KTH strain |  |
| TH29526 | *flhA9368*::tetRA(ΔAA137 of *flhA*) | TH27083 | tetRA 10718/10719 |
| TH29527 | *flhA9369*::tetRA(ΔAA82-AA92 of *flhA*) | TH27083 | tetRA 10710/10711 |
| TH29528 | *flhA9370*::tetRA(ΔAA242-AA249 of *flhA*) | TH27083 | tetRA 10722/10723 |
| TH29583 | *flhA7373*(T82V) | TH29527/pSim | LT2 10712/10713 |
| TH29584 | *flhA9375*(N89L) | TH29527/pSim | LT2 10714/10715 |
| TH29585 | *flhA9376*(N137) | TH29526/pSim | LT2 10727/10728 |
| TH29579 | *flhA9371*::tetRA(ΔAA201-AA219 of *flhA*) | TH27083/pSim | tetRA 10720/10721 |
| TH29580 | *flhB9372*::tetRA(ΔAA223-AA255 of *flhB)* | TH27083/pSim | tetRA 10725/10726 |
| TH29610 | *flhA9385*(S202A) | TH29579/pSim | LT2 10729/10730 |
| TH29611 | *flhA9386*(N219L) | TH29579/pSim | LT2 10731/9046 |
| TH29612 | *flhB9387*(R228A) | TH29580/pSim | LT2 10747/10748 |
| TH29613 | *flhB9388*(R249A) | TH29580/pSim | LT2 10749/10748  10750/10748 |
| TH29622 | *flhA9377*(T243V) | TH29528/pSim | LT2 10732/9361 |
| TH29623 | *flhA9378*(D249N) | TH29528/pSim | LT2 10742/10743 |
| TH29624 | *flhA9379*(S261A) | TH29523/pSim | LT2 10744/10745 |
| TH29625 | *flhA9380*(T93V) | TH29527/pSim | LT2 10717/10715 |
| TH29626 | *flhB9381*(R224A) | TH29580/pSim | LT2 10746/10748 |
| TH29627 | *flhB9382*(E233A) | TH29580/pSim | LT2 10749/10748  10750/10748 |
| TH29628 | *flhB9383*(R254A) | TH29580/pSim | LT2 10756/10753 |
| TH29657 | *flhA9389*(S92A) | TH29527/pSim | LT2 10716/10715 |
| TH29658 | *flhA9390*(T142V) | TH29526/pSim | Fill 10809/10810 |
| TH29659 | *flhB9391*(K241A) | TH29580/pSim | LT2 10768/10748  10750/10748 |
| TH29660 | *flhB9392*(K243A) | TH29580/pSim | LT2 10751/10748  10750/10748 |
| TH29661 | *flhB9393*(R245A) | TH29580/pSim | LT2 10752/10753  10754/10753 |
| TH29665 | *flhA9394*(Q254L) | TH29528/pSim | LT2 10807/10745 |
| TH29666 | *flhA9379*(S261A) *flhA9395*::tetRA(ΔAA137 of *flhA*) | TH29624/pSim | tetRA 10718/10719 |
| TH29667 | *flhA9386*(N219L) *flhA9396*::tetRA(ΔAA82 through AA94 of *flhA*) | TH29611/pSim | tetRA 10710/10711 |
| TH29668 | *flhA9379*(S261A) *flhA9397*::tetRA(DAA223-225 of *flhA*) | TH29624/pSim | tetRA 10720/10721 |
| TH29669 | *flhA9379*(S261A) *flhA9398*(N137L) | TH2966/pSim | Fill 10727/10728 |
| TH29670 | *flhA9376*(N137L) *flhA9399*(V305A) | TH29585 | Motile revertant |
| TH29671 | *flhA9376*(N137L) *flhA9400*(Δ(A107)) | TH29585 | Motile revertant |
| TH29672 | *flhA9376*(N137L) *flhA9401*(H101Q) | TH29585 | Motile revertant |
| TH29673 | *flhA9376*(N137L) *flhA9402*(A124V) | TH29585 | Motile revertant |
| TH29674 | *flhA9376*(N137L) *flhA9403*(L311P) | TH29585 | Motile revertant |
| TH29691 | *flhA9404*(S92D) | TH29527/pSim | LT2 10818/10715 |
| TH29692 | *flhA9405*(T246V) | TH29528/pSim | LT2 10741/9361 |
| TH29693 | *flhA9406*(T142V) | TH29526/pSim | LT2 10809/10810 |
| TH29694 | *flhA9407*(T142A) | TH29526/pSim | LT2 10814/10810 |
| TH29695 | *flhA9408*(N89S) | TH29527/pSim | LT2 10815/10715 |
| TH29696 | *flhA9409*(N89T) | TH29527/pSim | LT2 10815/10715 |
| TH29697 | *flhA9410*(S202A) *flhA9379*(S261A) | TH29668/pSim | LT2 10729/10730 |
| TH29698 | *flhA9411*(N219T) | TH29579/pSim | LT2 10816/9046 |
| TH29699 | *flhA9412*(D249S) | TH29528/pSim | Fill 10817/10743 |
| TH29700 | *flhA9413*(N89L) *flhA9386*(N219L) | TH29667/pSim | LT2 10714/10715 |
| TH29701 | *flhB9414*(Helix1 = R244S D226S R228S D229S E230S K232S E233S) | TH29580/pSim | LT2 10819/10748  10820/10748 |
| TH29702 | *flhB9415* (ΔAA241 through AA255 of *flhB*) | TH27083 | tetRA 10826/10726 |
| TH29733 | *flhA9416*(A253D) | TH29528/pSim | LT2 10825/10745 |
| TH29737 | *flhA9409*(N89T) *flhA9417*::tetRA (DAA201 through AA219 of *flhA*) | TH29696/pSim | tetRA 10720/10721 |
| TH29738 | *flhA9408*(N89S) *flhA9418*::tetRA (DAA242 through AA249 of *flhA*) | TH29695/pSim | tetRA 10722/10723 |
| TH29739 | *flhA9412*(D249S) *flhA9419*::tetRA (DAA81 through AA92 of *flhA*) | TH29699/pSim | tetRA 10710/10711 |
| TH29740 | *flhA9413*(N89L) *flhA9386*(N219L) *flhA9420*::tetRA (ΔAA137 of *flhA*) | TH29700/pSim | tetRA 10718/10719 |
| TH29741 | *flhA9410*(S202A) *flhA9379*(S261A) *flhA9421*::tetRA (ΔAA137 of *flhA*) | TH29697/pSim | tetRA 10718/10719 |
| TH29747 | *flhA9409*(N89T) *flhA9422*(N219T) | TH29737/pSim | LT2 10816/9046 |
| TH29748 | *flhA9408*(N89S) *flhA9423*(A253D) | TH29738/pSim | LT2 10825/10745 |
| TH29749 | *flhA9412*(D249S) *flhA9424*(S92D) | TH29739/pSim | LT2 10818/10715 |
| TH29750 | *flhA9413*(N89L) *flhA9386*(N219L) *flhA9425*(T142A) | TH29740/pSim | Fill 10814/10810 |
| TH29751 | *flhA9410*(S202A) *flhA9379*(S261A) *flhA9426*(T142V) | TH29741/pSim | Fill 10814/10810 |
| TH29752 | *flhB9427*(helix1: R224A/D226A/R228A/D229A/E230A/K232A/E233A) | TH29580/pSim | LT2 10842/10748  10843/10748 |
| TH29753 | *flhB9428*(helix1: R224S/R228S/K232S) | TH29580/pSim | LT2 10846/10748  10847/10748 |
| TH29754 | *flhB9429*(helix2: K241S/K243S/R245S/R249S/R254S/R255S) | TH29702/pSim | Fill 10821/10822  PCR 10823/10824 |
| TH29755 | *flhB9430*(helix2: K241A/K243A/R245A/R249A/R254A/R255A) | TH29702/pSim | Fill 10848/10849  PCR 10823/10824 |
| TH29756 | *flhB9431*(helix2: K241S/R245S/R249S) | TH29702/pSim | Fill 10850/10851  PCR 10823/10824 |
| TH29786 | *flhA9386*(N219L) flhA9437(S92A T93A) | TH29667/pSim | LT2 10808/10715 |
| TH29787 | *flhB9438*(helix2: K243S/K254S/R255S) | TH29702/pSim | LT2 10852/10853  10823/10824 |
| TH29816 | *flhB9444*(helix 1: D226S D229S E230S) | TH29702/pSim | LT2 10844b/10748  10845/10748 |
| TH29823 | *flhB9445*(helix 1: D226S D229S E230S E233S) | TH29702/pSim | Fill 10844/10748  PCR 10845/10748 |

**Table S2. List of primers:**

| **Name** | **Sequence (5’ to 3’)** |
| --- | --- |
| 9046-flhA-aa198-201+43bp-fw | acggcgttcggaagtcac |
| 9361-fillin- flhatm6 | atcggtactaacgcgggt |
| 10710-flha246tetr | tctggattttgccgcgtttccgaccatcctgctgtttaccttaagacccactttcacatt |
| 10711-flha279-teta | ccgccgccgcgccggtatgcccttccatcaaaataatgcgctaagcacttgtctcctg |
| 10712-flha-t82vfw | tctggattttgccgcgtttccgaccatcctgctgtttaccgtgctactgcgtctggcgctta |
| 10713-flha-bp343 | cgaacgcctccactaccttt |
| 10714-flha-n89lrv | ccgccgccgcgccggtatgcccttccatcaaaataatgcgcgttgaggcaacgagaagcgccagacgcagtag |
| 10715-flha-2 | gatgtttacccagcgtactct |
| 10716-flha-s92a-rev | ccgccgccgcgccggtatgcccttccatcaaaataatgcgcgttgcggcaacgttaagcgccag |
| 10717-flha-t93vrev | ccgccgccgcgccggtatgcccttccatcaaaataatgcgcactgaggcaacgttaagcgc |
| 10718-flha411tetr | cgcgatcgggattgtggtttttatcatcctcgtcattatcttaagacccactttcacatt |
| 10719-flha 411teta | gacttcggcaatacgtccggcgcctttggtgatgaccatgaactaagcacttgtctcctg |
| 10720-flha603-tetr | cactcaggaagccgacttctacggctcgatggacggggcattaagacccactttcacatt |
| 10721-flha-657teta | cattccgtgctgcaatacgccgaccagcagcccgccgaccacctaagcacttgtctcctg |
| 10722-flha729-tetr | gcagcacggaatgagcataggcagcgcggcggaaagctacttaagacccactttcacatt |
| 10723-flha-747-teta | ccgtggagataaccagcgccgggatctgggcgaccaggccctaagcacttgtctcctg |
| 10725-flhb-672bp-tetr | tttccagatctttagccacctgaaaaaattacgcatgtcgttaagacccactttcacatt |
| 10726-flhb-767bp-teta | tagtgacaatgacgtccgctttcggcacatcttccatcatctaagcacttgtctcctg |
| 10727-flha-n411lfw | cgcgatcgggattgtggtttttatcatcctcgtcattatcctcttcatggtcatcacca |
| 10728-flha-n411l-rv | gacttcggcaatacgtccggcgcctttggtgatgaccatgaag |
| 10729-flha-s202afw | cactcaggaagccgacttctacggctcgatggacggggcagctaaatttgtacgcggcgac |
| 10730-flha202-rev | tgcctatgctcattccgt |
| 10731-flha-n219lrev | ttccgtgctgcaatacgccgaccagcagcccgccgaccacgagaatgaccatgatgagaata |
| 10732-flha-t243vfw | gcagcacggaatgagcataggcagcgcggcggaaagctacgtcctgctgaccattggcgac |
| 10741-flha-t246v | gcagcacggaatgagcataggcagcgcggcggaaagctacaccctgctggtcattggcgacggcctggtc |
| 10742-flha-d249n | ccgtggagataaccagcgccgggatctgggcgaccaggccgttgccaatggtcagcagggt |
| 10743-flha-d249n-fill | gcagcacggaatgagcataggcagcgcggcggaaagctacaccctgctgaccattg |
| 10744-flha-s261a | gatcggtactaacgcgggtcacaatgacgcccgccgccgtggcgataaccagcgccgggat |
| 10745-flha-mid | gcgtattgcagcacggaatg |
| 10746-flhb-r224afw | tttccagatctttagccacctgaaaaaattacgcatgtcggcgcaggacattcgcgacgaa |
| 10747-flhb-r228afw | tttccagatctttagccacctgaaaaaattacgcatgtcgcggcaggacattgccgacgaatttaaagagagcgaa |
| 10748-flhb-rev | gtcgggttagtgacaatgacg |
| 10749-flhb-e233a-fw | cggcaggacattcgcgacgaatttaaagcgagcgaaggcgatccgcat |
| 10750-flhb-fill | tttccagatctttagccacctgaaaaaattacgcatgtcgcggcaggacattcgcgac |
| 10751-flhb-k243a-fw | cggcaggacattcgcgacgaatttaaagagagcgaaggcgatccgcatgttaagggcgcaattcgccagatgcaacg |
| 10752-flhbr245a-rev | ttcggcacatcttccatcatgcggcgctgcgcggcggcgcgttgcatctgggcaattttgcccttaacatgc |
| 10753-flhb-241-fill | ggtgggatttgacgtgtt |
| 10754-flhb245-fill | gggttagtgacaatgacgtccgctttcggcacatcttccatc |
| 10756-flhb-r254arv | tagtgacaatgacgtccgctttcggcacatcttccatcatgcgggcctgcgcggcggcgcgttg |
| 10768-flhb-k241a-fw | cggcaggacattcgcgacgaatttaaagagagcgaaggcgatccgcatgttgcgggcaaaattcgccagatg |
| 10807-flha-q254l-rv | caatgacgcccgccgccgtggagataaccagcgccgggatcagggcgaccaggccgtcgc |
| 10808-flha-s92a-t93v | cgccgccgcgccggtatgcccttccatcaaaataatgcgcactgcggcaacgttaagcgccag |
| 10809-fill-flha142v-rev | caaagcgcgcgccgacttcggcaatacgtccggcgcctttgacgatgaccatgaagttgat |
| 10810-flha-t142vfw | cgcgatcgggattgtggtttttatcatcctcgtcattatcaacttcatggtcatc |
| 10814-fill-flha142a-rev | caaagcgcgcgccgacttcggcaatacgtccggcgcctttggcgatgaccatgaagttgat |
| 10815-flha-n89t-lrv | ccgccgccgcgccggtatgcccttccatcaaaataatgcgcgttgaggcaacngwaagcgccagacgcagtag |
| 10816-flha-n219trev | ttccgtgctgcaatacgccgaccagcagcccgccgaccacggtaatgaccatgatgagaata |
| 10817-flha-d249s | ccgtggagataaccagcgccgggatctgggcgaccaggccgctgccaatggtcagcagggt |
| 10818-flha-s92d-rev | ccgccgccgcgccggtatgcccttccatcaaaataatgcgcgtatcggcaacgttaagcgccag |
| 10819-flhb-7ser-helix1-pcr1 | tcgcagagcatttctagctcgtttagtagcagcgaaggcgatccgcat |
| 10820-flhb-7ser-helix1-pcr2 | tttccagatctttagccacctgaaaaaattacgcatgtcgtcgcagagcatttctagc |
| 10821-flhb-helix2-ser | cggcacatcttccatcatactgctctgcgcggcggcactttgcatctggctaatactgccgc |
| 10822-flhb-helix2ser-fill | cgcgacgaatttaaagagagcgaaggcgatccgcatgttagcggcagtattagccaga |
| 10823-flhb-fill | tagtgacaatgacgtccgctttcggcacatcttccatcat |
| 10824-flhb-255-rv | tcgcgacgaatttaaagagag |
| 10825-flha253d-rev | tgacgcccgccgccgtggagataaccagcgccgggatctggtcgaccaggccgtcgccaa |
| 10826-flhb-241-tetr | cgcgacgaatttaaagagagcgaaggcgatccgcatgttattaagacccactttcacatt |
| 10842-flhb-7alagcn-helix1-pcr1 | ttacgcatgtcggcgcaggccattgccgccgcatttgcagcgagcgaaggcgatccgcat |
| 10843-flhb-7ala-helix2-pcr2 | tttccagatctttagccacctgaaaaaattacgcatgtcggcgcag |
| 10844-flhb-de-helix1-pcr1 | ttacgcatgtcgcggcagagcattcgcagctcgtttaaaagcagcgaaggcgatccgcat |
| 10844b-flhb-de-helix1-pcr1 | ttacgcatgtcgcggcagagcattcgcagctcgtttaaagagagcgaaggcgatccg |
| 10845-flhb-helix1-fill-wt | tttccagatctttagccacctgaaaaaattacgcatgtcgcggcag |
| 10846-flhb-helix1kr-pcr1 | atgtcgtcgcaggacatttctgacgaatttagtgagagcgaaggcgatccg |
| 10847-flhb-h1-kr-fill | tttccagatctttagccacctgaaaaaattacgcatgtcgtcgcaggacatt |
| 10848-flhb-helix2-ala | cggcacatcttccatcatggctgcctgcgcggcggcggcttgcatctgggcaattgcgccggc |
| 10849-flhb-helix2-ala-fill | cgcgacgaatttaaagagagcgaaggcgatccgcatgttgccggcgcaattgcccag |
| 10850-flhb-helix2-241-245-249 | cggcacatcttccatcatgcggcgctgcgcggcggcactttgcatctggctaattttgccgc |
| 10851-flhb-241-245ser-fill | cgcgacgaatttaaagagagcgaaggcgatccgcatgttagcggcaaaattagccaga |
| 10852-flhb-helix2-243-254-255-s | cggcacatcttccatcatactgctctgcgcggcggcgcgttgcatctggcgaatactgcc |
| 10853-flhb-helix2-243-254-255-s-fill | cgcgacgaatttaaagagagcgaaggcgatccgcatgttaagggcagtattcgccagatg |

**Table S3. Cryo-EM data collection, refinement and validation statistics**

|  | Export Gate with FlhB  (EMD-77586)  (PDB 36HU) | FlhA_TM_  (EMD-77588)  (PDB 36HW) | Export Apparatus  (EMD-77603)  (PDB 36IS) |
| --- | --- | --- | --- |
| **Data collection and processing** |  |  |  |
| Magnification | 165,000 | 165,000 | 165,000 |
| Voltage (kV) | 300 | 300 | 300 |
| Electron exposure (e–/Å^2^) | 58.5 | 58.5 | 58.5 |
| Defocus range (μm) | -2.5 to -1.0 | -2.5 to -0.5 | -2.5 to -0.5 |
| Pixel size (Å) | 0.732 | 0.732 | 0.732 |
| Symmetry imposed | C1 | C9 | C1 |
| Initial particle images (no.) | 355242 | 355242 | 355242 |
| Final particle images (no.) | 151443 | 19350 | 58082 |
| Map resolution (Å)  FSC threshold | 2.5  0.143 | 3.6  0.143 | 4.3  0.143 |
| **Refinement** |  |  |  |
| Model resolution (Å)  FSC threshold | 2.5  0.143 | 3.6  0.143 | 4.3  0.143 |
| Map sharpening B factor (Å^2^) | -58 | -39 | -77 |
| Model composition  Non-hydrogen atoms  Protein residues | 14357  1865 | 20781  2799 | 35138  4664 |
| B factors (Å^2^)  Protein | 38.8 | 81.9 | 149.1 |
| R.m.s. deviations  Bond lengths (Å)  Bond angles (°) | 0.004  0.593 | 0.010  0.996 | 0.002  0.614 |
| Validation  MolProbity score  Clashscore  Poor rotamers (%) | 1.5  6.23  0.19 | 2.3  18.2  0.41 | 1.9  9.1  0.08 |
| Ramachandran plot  Favored (%)  Allowed (%)  Disallowed (%) | 97.1  2.8  0.1 | 89.6  10.4  0.0 | 94.7  5.3  0.0 |
| CC (mask) | 0.83 | 0.87 | 0.82 |
